# Transient spectral events in resting state MEG predict individual time-frequency task responses

**DOI:** 10.1101/419374

**Authors:** R Becker, D Vidaurre, AJ Quinn, R Abeysuriya, O Parker Jones, S Jbabdi, MW Woolrich

**Author notes:** Correspondence to: Robert Becker.

## Abstract

Even in response to apparently simple tasks such as hand moving, human brain activity shows remarkable inter-subject variability. Presumably, this variability reflects genuine behavioural or functional variability. Recently, spatial variability of resting-state features in fMRI - specifically connectivity - has been shown to explain (spatial) task-response variability. Such a link, however, is still missing for M/EEG data and its spectrally rich structure. At the same time, it has recently been shown that task responses in M/EEG can be well represented using transient spectral events bursting at fast time scales. Here, we show that individual differences in the spatio-spectral structure of M/EEG task responses, can, to a reasonable degree, be predicted from individual differences in transient spectral events identified at rest. In a MEG dataset of diverse task conditions (including motor responses, working memory and language comprehension tasks) and resting-state sessions for each subject (n = 89), we used Hidden-Markov-Modelling to identify transient spectral events as a feature set to learn the mapping of space-time-frequency content from rest to task. Resulting trial-averaged, subject-specific task-response predictions were then compared with the actual task responses in left-out subjects. All task conditions were predicted significantly above chance. Furthermore, we observed a systematic relationship between genetic similarity (e.g. unrelated subjects vs. twins) and predictability. These findings support the idea that subject-specific transient spectral events in resting-state neural activity are linked to, and predictive of, subject-specific trial-averaged task responses in a wide range of experimental conditions.

## 1. Introduction

Human non-invasive neuroimaging data is characterized by high inter-subject variability. Even for simple tasks such as moving a hand or seeing a well-defined visual pattern, the specific responses elicited in different subjects can be heterogeneous in terms of spatial location or extent, as well as magnitude, timing and oscillatory content. The origin of this variability is not clear, but there is increasing evidence that it reflects intrinsic inter-individual differences in resting-state activity. Support for this hypothesis has been demonstrated recently using human functional magnetic resonance imaging (fMRI) data, where spatial activation maps for a number of different tasks (motor, sensory, working memory) were reliably predicted from connectivity profiles derived from resting state data (Tavor et al., 2016). Other fMRI studies examining functional connectivity patterns showed that these patterns - both in rest and task - can serve as unique, subject-specific signatures of brain activity, also known as neuronal ‘fingerprinting’ (Finn et al., 2015) indicating another potential link between rest and task brain activity. However, these studies and others, focusing on rest-task relationships (e.g. Cole et al., 2016), were being undertaken in the domain of fMRI.

However, a common criticism of fMRI is that it represents a rather indirect measure of brain activity. This leads to the question whether the same relationship or predictability holds for more direct measures of brain activity such as magnetoencephalography (MEG) or electroencephalograpy (EEG). M/EEG can capture the oscillatory and synchronized activity of neuronal populations and - unlike fMRI - can resolve brain activity at a temporal resolution down to milliseconds, reaching the temporal scale at which important aspects of cognition, and the neural dynamics that are tied to these processes, arise. Thus, the natural question arises whether features of M/EEG task responses can be predicted from rest as well. There is already a large body of work focusing on the link between rest and task processing in M/EEG. Features of the most prominent rhythms, such as alpha or beta oscillations, in human resting state M/EEG data are known to be functionally relevant and have considerable cross-subject variability (Klimesch, 1999). As in fMRI, the M/EEG literature has already demonstrated both the variability of transient electrophysiological features at rest and also during task-processing and their functional relevance. For example, resting state features in EEG have been shown to impact on task responses (Becker, Ritter, & Villringer, 2008; Mazaheri & Jensen, 2008; Nikulin et al., 2007), relate to perceptual and cognitive performance (Busch, Dubois, & VanRullen, 2009), (Mathewson, Gratton, Fabiani, Beck, & Ro, 2009). On the other hand, ongoing activity in the alpha or beta frequency range is also highly variable during task processing and shows functional relevance during working memory tasks (Klimesch, 1997), observation of movements (Pineda et al., 2005) as well during effortful speech comprehension (Becker, Pefkou, Michel, & Hervais-Adelman, 2013; Obleser & Weisz, 2012). The link between ongoing and task activity has been demonstrated both intra-individually (i.e. on a trial-by-trial level) and inter-individually (i.e. on a subject-by-subject level), and such links seems to exist beyond specific sensory and cognitive domains and have been demonstrated also invasively in animal brain activity (e.g. (Arieli, Sterkin, Grinvald, & Aertsen, 1996)). While all these studies showed evidence of some link or interaction between rest and task activity in neuronal activity, a more direct link – i.e. predicting individualised task responses from spontaneous neuronal activity - is still missing for M/EEG.

In this study, building on the work of Tavor et al. (2016) that focused on fMRI spatial variability, we aim to predict cross-subject variability of task MEG time-frequency responses using spatio-spectral dynamics as derived from resting MEG data. Tavor et al. (2016) succeeded in predicting task fMRI activation spatial maps using spatial (i.e. connectivity) properties derived from resting fMRI data by using modes that corresponded to spatially-localised sub-portions of the connectivity profiles extracted from the individual static functional connectivity network matrices. Here, for predicting spectrally rich M/EEG task responses, and since we are bringing in the spectro-temporal dimension, we require a method that can reliably extract the relevant spectral subject-specific properties (or modes) from rest data in an unsupervised fashion while accommodating the dynamics contained in the data.

There are a number of possible approaches for extracting the required dynamic spatio-spectral modes from resting M/EEG data; including ICA and sliding-window estimates of functional connectivity (Brookes et al., 2011; Hipp, Hawellek, Corbetta, Siegel, & Engel, 2012). However, it has recently been shown that task responses in M/EEG can be well represented using transient spectral events bursting at fast time scales (van Ede et al., 2018, Shin et al., 2017; Vidaurre et al., 2016; Zich et al., 2018). This work has revealed that sustained oscillations in task data observed after trial-averaging may actually correspond to the temporal smearing of responses bursting at fast timescales with variable timing. We therefore hypothesised that subject-specific transient spectral events in resting-state neural activity might be predictive of subject-specific trial-averaged task responses, in a wide range of experimental conditions.

Here, we looked to test this idea by using the approach of Hidden-Markov-Modeling (HMM) to identify fast transient spectral events, since the HMM has previously been shown to be capable of extracting events with rich spectral profiles in both rest and task MEG data. For example, the HMM has proven useful in resting MEG data for identifying fast transient events with distinct multi-regional power amplitudes and amplitude correlations (Baker et al., 2014), or with distinct multi-regional spectral and cross-spectral (e.g. coherence) properties (Vidaurre et al., 2018); and in task MEG data for capturing task-modulations of spectral properties (Vidaurre et al., 2016; Zich et al., 2018).

We sought to predict between-subject variability in the time-frequency responses in a number of different tasks using resting state data. For this we use data from the ‘Human Connectome Project’ (HCP), a consortium of several research institutes that has collected data of a relatively large number of MEG subjects and incorporates both resting state data and a range of task data, including motor movements, and cognitive tasks involving working-memory and language comprehension (Van Essen et al., 2013).

## 2. Methods

### 2.1. Subjects and data

The data used here are human non-invasive resting state and task magnetoencephalography (MEG) data publicly available from the Human Connectome Project (HCP) consortium (Van Essen et al., 2013, Larson-Prior et al., 2013), acquired on a Magnes 3600 MEG (4D NeuroImaging, San Diego, USA) with 248 magnetometers. The resting state data consist of 89 subjects (mean 28.7 years, range 22-35, 41 f / 48 m, acquired in 3 subsequent sessions, lasting 6 minutes each). Task data were available for a subset of these subjects, with 2 sessions per task. Each task has a similar or identical experimental design to the corresponding task acquired during fMRI imaging. The tasks acquired in the MEG include a motor task condition - where hands or feet had to be moved paced by an external cue (every 1.2s), a working memory (WM) task - where people had to remember the occurrence a of n-back previously shown item (with n=0 and 2) with the items being either tools or faces and finally, a third task group - involving language comprehension - where in one condition, subjects had to listen to a number of sentences (making up a complete story) and then answer questions regarding that story, and in another condition subjects had to solve math problems (Larson-Prior et al., 2013). Data were segmented to the onset of EMG (motor task), the non-target item (WM task), or to the onset of a sentence (language task). For each of the task groups, overlapping but not identical subsets of the total pool of resting state subjects was available (motor task n=56, WM task n=70, language comprehension n=72).

We analysed 10 different task conditions in total across these 3 main task groups – 4 for the motor task (right hand, left hand, right foot, left foot), 4 for the WM task (0-back face items, 0-back tool items, 2-back face items, 2-back tool items, all being part of the non-target condition which also required a motor response), and 2 task conditions within the language condition - sentences vs. math problems.

### 2.2. Preprocessing

#### 2.2.1. Source estimation and parcellation

For each subject, the MEG data were acquired in a single continuous run comprising both rest and task. We used the MEG data from the HCP database denominated as ‘preprocessed’ as starting point. At this level of preprocessing, removal of artefactual independent components, bad samples and channels had been already performed (see Larson-Prior et al., 2013). Following these steps, the data were subject to bandpass filtering (1-48Hz, Butterworth) and LCMV beamforming (using beamforming routines from the Matlab based Fieldtrip toolbox (Oostenveld, Fries, Maris, & Schoffelen, 2011)), resulting in 5798 virtual source voxels (with 8mm grid resolution) and down-sampled to 200 Hz. In order to reduce dimensionality of the analysed data, we used a custom parcellation of 76 parcels covering the whole brain, extracting the first principal component (PC) across all time-courses within each parcel. The parcellation was created in a way such that each first PC explained about 60% of the variance across all voxels within each parcel by starting with 2 large parcels covering each hemisphere and then subsequently splitting these parcels into smaller ones). The parcellation was based – analogous to all other preprocessing steps – on the resting state only.

Note that all resting state runs for a subject were acquired in a single session. As a result, we concatenated the resting state runs for a subject, and applied a single beamformer, parcel time-course extraction and spatial leakage reduction. Then, the transformations (learnt only from the rest data for a subject) are applied to the task data runs for any subjects we are looking to predict. This ensures maximum consistency of within-subject pre-processing for all sessions, without having knowledge of or being biased by any task data information of these subjects.

#### 2.2.2. Spatial leakage reduction

For spatial leakage reduction, we chose the multi-variate orthogonalisation approach as described in (Colclough, Brookes, Smith, & Woolrich, 2015), using the ‘closest’ implementation).

#### 2.2.3. Task epoching

Task data was segmented into epochs depending on the event of interest: For the motor task, data were time-locked to the onset of the electromyogram (EMG), for the WM task, epochs were locked to the visual onset of the (non-target) item and for the language comprehension task the beginning of the sentence or maths problem was the time-locking event. To ensure conformity of the concatenated data sets, both resting state data and task data were normalized to zero mean and unit variance (performed per subject and parcel). Motor task epochs were segmented from −1.1 to 1.1secs, working memory task data and language task data from −1.1 to 2.2 secs. Baseline correction was performed from −0.5 to - 0.2s.

### 2.3. Conventional Estimate of Time-Frequency Task Responses

We performed conventional wavelet (WL) time-frequency analysis (7 cycles, Morlet mother wavelet) for the segmented task data. The resulting WL-based task responses serve as comparison with the HMM-based (regularised) task responses at the group level. The WL based task responses were baseline corrected in a pre-stimulus time window (−0.5s to −0.2s for all task conditions). If not specified otherwise, general custom Matlab scripts were used (Matlab R2016b, Natick, USA) and the in-house OHBA Software Library (OSL), which is built on Fieldtrip and SPM, and is available at https://github.com/OHBA-analysis/osl-core).

### 2.4. Prediction of Subject-Specific Time-Frequency Task Responses using Resting Data

After pre-processing, resting state parcel time-courses were analysed one parcel at a time using the pipeline outlined in **Fig. 1**. In **Supplementary Information (SI) Fig. 1**, the approach is illustrated in more detail (including usage of the HMM). In general, the pipeline has the following steps: In the training step, we first identify group and individual state time courses and their (spectral) signatures in rest. Note that each HMM state corresponds to a spectral event of certain type, i.e. with a distinct spectral profile. We then estimate - in task - individual state time courses by fitting the rest states to the task data. In the general framework (**Fig. 1**), in the prediction step we then predict – in the unseen subject – its own task response by combining their individual resting state signatures with the observed state time course for the group.

**Figure 1.**
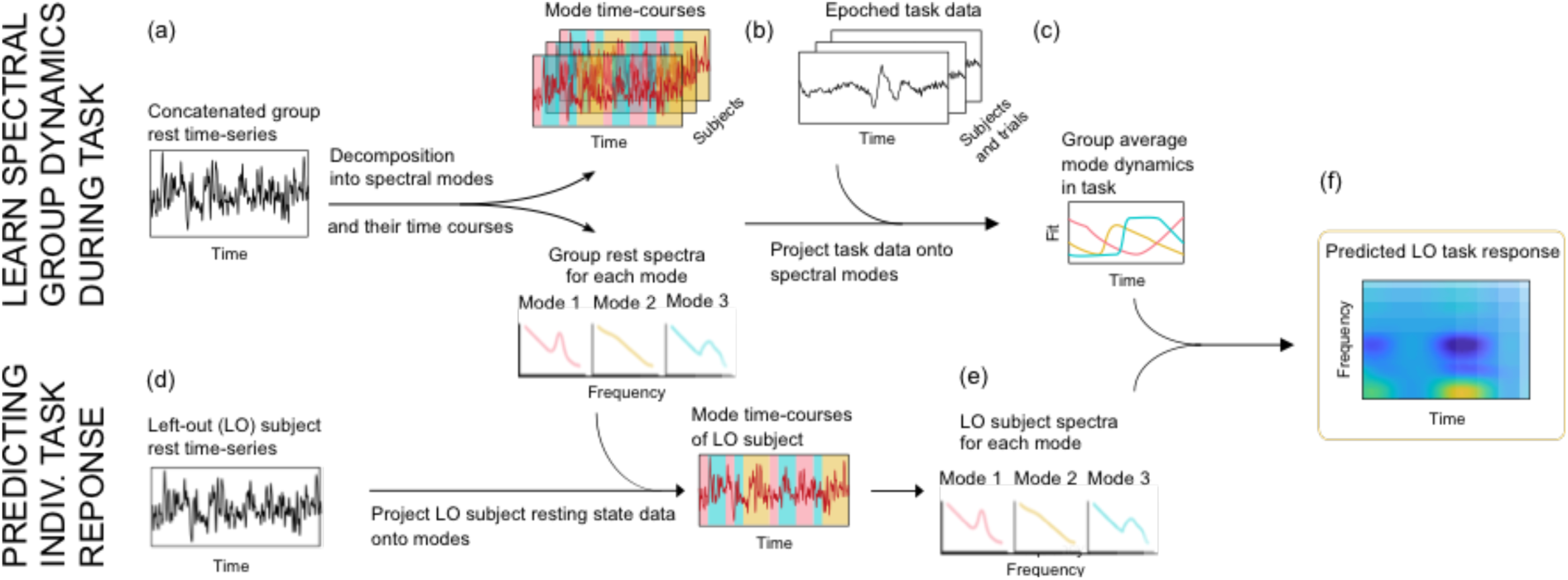
An overview of the approach shown for one task condition, and which is carried out separately on all parcels. **a-c. Training stage**. (**a**) Spectral modes are identified on the entire group resting state data (e.g. by using HMM, where modes correspond to different types of spectral events represented by the HMM states), resulting in the group-averaged rest spectra for each mode. (**b**). The epoched task data for all subjects and trials is projected onto the group-averaged mode spectra at rest, resulting in group-averaged task-locked dynamics for each mode (**c**). **d-f. Prediction of the task-response** for the left-out (LO) subject. (**d**) The LO subject’s resting state activity, from panel (a), is projected onto the group mode spectra to create subject-specific state-time courses and subsequently, subject-specific spectral modes (**e**). Finally, the group-averaged task-locked mode dynamics, from panel (c), and the LO subject-specific mode spectra at rest, from panel (e), are combined to create a LO subject-specific prediction of the time-frequency task response (**f**). This is a simplification of our approach, a more detailed schematic, specific to the use of the HMM, is depicted in **Supplementary Fig. 1**.

#### 2.4.1. Training

As illustrated in **Fig. 1a-c** (and in more detail in **SI Fig. 1a-c**) we train as a first step (**SI Fig. 1a**) an autoregressive Hidden-Markov-Model (HMM-AR, with 4 hidden states and an autoregressive (AR) observation model with order 5) to identify states representing different types of spectral events in the ***group resting*** state data (concatenated over subjects, n=89). We used the HMM toolbox, available at https://github.com/OHBA-analysis/HMM-MAR. Hidden-Markov Modelling is a stochastic modelling approach that considers the data as being generated from a discrete sequence of hidden underlying states. The HMM infers both the characteristics of the states and their time courses simultaneously. Within the HMM, the observation model (sometimes referred to as the emission distribution) describes, for each state, the distribution explaining the data when that state is active. In our case the observation model is a univariate autoregressive (AR) model of order 5, meaning that the hidden states basically correspond to a spectral event of certain type, i.e. with a distinct spectral profile, and that each visit to a state corresponds to a distinct spectral event of that type. We refer to this approach as an HMM-AR - for further details of the approach see Vidaurre et al., (2016), where the more general case of a multivariate AR observation model is discussed. Here, we infer separate HMM-AR parameters for each brain region (or parcel), resulting in the state time-courses (indicating the probability of the state being active at each time point) per state and subject, and the group-averaged state spectra at rest (i.e. the state observation models or the spectral profiles of the different spectral event types). The group-averaged state spectral properties of each spectral event type or state at rest are computed as described in section 2.4.3, “State Spectra Estimation” using a multi-taper approach.

Subsequently (**SI Fig. 1b**), we infer the HMM-AR on the epoched task data for all subjects and trials, but, crucially, with the observation model held fixed to the pre-estimated group-averaged state spectra at rest (i.e. from the previous step). The resulting state time-courses are trial-averaged to give the subject-specific task-locked dynamics for each state; we refer to these average state time courses as “occupancies”, indicating the relative amount of time spent in that state at each time point (or alternatively, the rate of occurrence of the spectral event types defined by that state).

Finally (**SI Fig. 1c**), as part of a leave-one-out cross validation scheme, we learn a linear mapping from each individual’s average rest occupancy to their own actual task occupancy at each time point within the task window, leaving out one subject. This will be used to improve the prediction of the subject-specific state occupancies in task (i.e. the subject-specific task-locked dynamics for each state). The most straightforward way to predict the subject-specific state occupancies in task would be to simply set them to be the same as the group-averaged occupancies in task (as in **Fig. 1**). However, this would not take into account the extent to which individuals tend to spend different amounts of time in each state; and assuming that there is a relationship between the amount of time spent in a state at rest and in the task, taking this into account will improve our prediction of the subject-specific task responses. This linear mapping (see next methods section “Prediction” for details) is learned separately at each time-point within trial, by using linear regression across all subjects between the subject-specific state occupancies in rest and the subject-specific states occupancies in task at the time-point in question. As a result, we obtain regression coefficients that vary as a function of time within the trial (**SI Fig. 1c**).

#### 2.4.2 Prediction

This is illustrated in **Fig. 1d-f** and in more detail in **SI Fig. 1d-f**. In the prediction stage, we will make use of the trained model, and apply it to a new, left-out (LO) subject to predict their unseen time-frequency task response. As with the training, this is carried out one parcel at a time.

First, the HMM-AR is fit to the LO subject’s rest data, but with the observation model held fixed to the group-averaged state spectra at rest (from **SI Fig. 1a**), resulting in the LO subject’s state time-courses at rest (**SI Fig. 1d**). These state time-courses are used to obtain LO subject’s state occupancies and state spectra at rest. The state-specific spectra are estimated by the method described in section 2.4.3, “State Spectra Estimation”.

Next, we predict the LO subject’s state occupancies in task (i.e. the LO subject’s task-locked dynamics for each state). As mentioned previously, the most straightforward thing to do, would be to simply set the predicted LO subject’s state occupancies in the task to be the same as the group-averaged state occupancies in the task (from **SI Fig. 1b**). However, we instead take into account the extent to which individuals tend to spend different amounts of time in each state, by assuming that that is a related quantity between rest and task. For each trial time-point, this is done by taking each subject’s average state occupancies at rest (SOR, collapsed over time m= 1…. m, for all n subjects) :

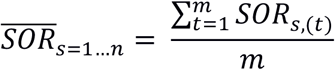

Having obtained the rest state occupancies (SOR) for all subjects, we perform a linear regression on all subjects except the left out subject (n being the index of the left out subject) to find the linear relationship between the state occupancy in a task at time t (SOT) and the average state occupancy at rest (SOR) across all subjects (except LO subject, n). This linear relationship is identified for each state (s), parcel (p), task (k) and time (t)):

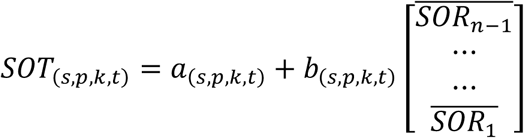

The obtained coefficients *a* and *b* are then used to predict the LO subject’s state occupancies in task (**SI Fig. 1e**), (again, separately for each state, parcel, task and time):

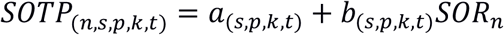

The above leave-one-out pipeline (see also **Supp Fig. 1**) is repeated for all n subjects, so that every subject has a predicted SOT for all times, tasks, and parcels (without having seen its own task data).

Finally, the predicted LO subject’s state occupancies in task (time x states) and the LO subject’s state spectra at rest (frequency x states) are combined (by matrix multiplication), to give a prediction of the LO subject’s time-frequency task response (time x frequency) (**SI Fig. 1f**). These are then baseline corrected in the same way as the conventionally produced time-frequency responses (see section 2.3., “Conventional Estimate of Time-Frequency Task Responses”). Henceforth, these will be referred to as the ‘***predicted task responses’***.

#### 2.4.3. State spectra estimation

Once the state time-courses are defined, we refine the estimate of their spectral profiles. Recall that these spectral profiles capture the nature of the type of the spectral events that each state represents. While this can be done in a parametric way, i.e. using the parameters obtained by the observation model (i.e. the AR coefficients in our case, e.g. see (Vidaurre et al., 2016)), we choose here to do this non-parametrically by means of a multi-taper spectral approach. Analogous to (Vidaurre et al., 2016), state-wise spectral power is computed as follows:

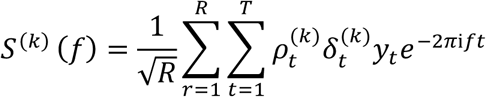

With

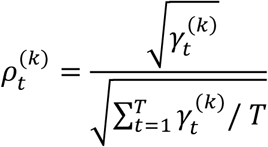

where *ρ* is weighting the spectral power at time point t by how much a point is represented by state k. These derived state spectra will serve as input into the reconstruction and prediction step of our approach pipeline (see **Fig. 1 and SI Fig. 1**, see Methods section 2.4.1, “Training”). State spectra are computed once per subject and parcel. The group-average state spectra will be used to created group-average reconstructed time-frequency responses that will be compared against the conventional wavelet-based time-frequency responses.

#### 2.4.4 Validation of prediction approach

In the validation step, we assess the quality of the prediction by comparing the LO subject’s time-frequency predicted task response (from **SI Fig. 1F**) to their actual task response. The HMM-AR limits the dimensionality of the predicted task response (because the AR model has only 5 parameters, and because there are only 4 states) so it is by design not capable of capturing all features of the actual task response. To account for this in the comparison, we therefore compare the predicted response to the ‘HMM regularized’ estimate of the LO subject’s task response. This was computed by fitting the HMM-AR to the LO subject’s actual task data, but while holding the observation models fixed to be the group-averaged state spectra at rest (as we do in the prediction). This results in state occupancies and the associated state spectra in the task for the LO subject which are then combined (by means of matrix multiplication), to give ‘HMM regularized’ time-frequency task responses for the LO subject. These are baseline corrected in the same way as the conventionally produced time-frequency responses (see section 2.3., “Conventional Estimate of Time-Frequency Task Responses”). Henceforth, these HMM regularized TF estimates will be referred to as the ‘***actual task*** responses’, and the conventional (wavelet-based) time-frequency task responses as ‘***WL based task responses’***.

Finally, to assess the quality of the prediction, a linear correlation analysis is used to compare the predicted and actual task response for the LO subject. Before the correlation, predicted task responses (i.e. 2D time-frequency maps) are concatenated over all parcels for each LO subject. If the prediction is performing sufficiently well, then this predicted task response should be more similar to the actual task response of the LO subject than to the actual task responses of any other subject. In a second step, correlation coefficients are normalized (demeaned, with unit variance) over rows and columns. In order to determine the statistical significance of this effect, a two-sample t-test is performed testing whether diagonal and off-diagonal samples come from the same distribution.

### 2.5. Sources of variability of prediction performance across tasks and subjects

Next, we examine whether there is any systematic variation in how well we can predict different tasks and different subjects. Specifically, we investigate the relationship between prediction performance and a measure of the ‘stability’ of each subject’s first-level task. Prediction performance was quantified using the average diagonal value of the correlation matrix (see **Fig. 5**), and the task stability was quantified (within subjects) using the resulting t-value over all trials for a time window of interest from 0.1s to 0.5s post-stimulus, frequency range from 8-26 Hz (testing the difference from zero), and for a representative parcel-of-interest for each task, as shown in **Fig. 3**. For the motor task, we chose a parcel corresponding to the contra-lateral motor area, for the working memory task we chose a parcel in proximity to visual cortex, and for the language task a parcel near the auditory cortex was chosen. These measures are computed for every task and subject.

**Figure 2.**
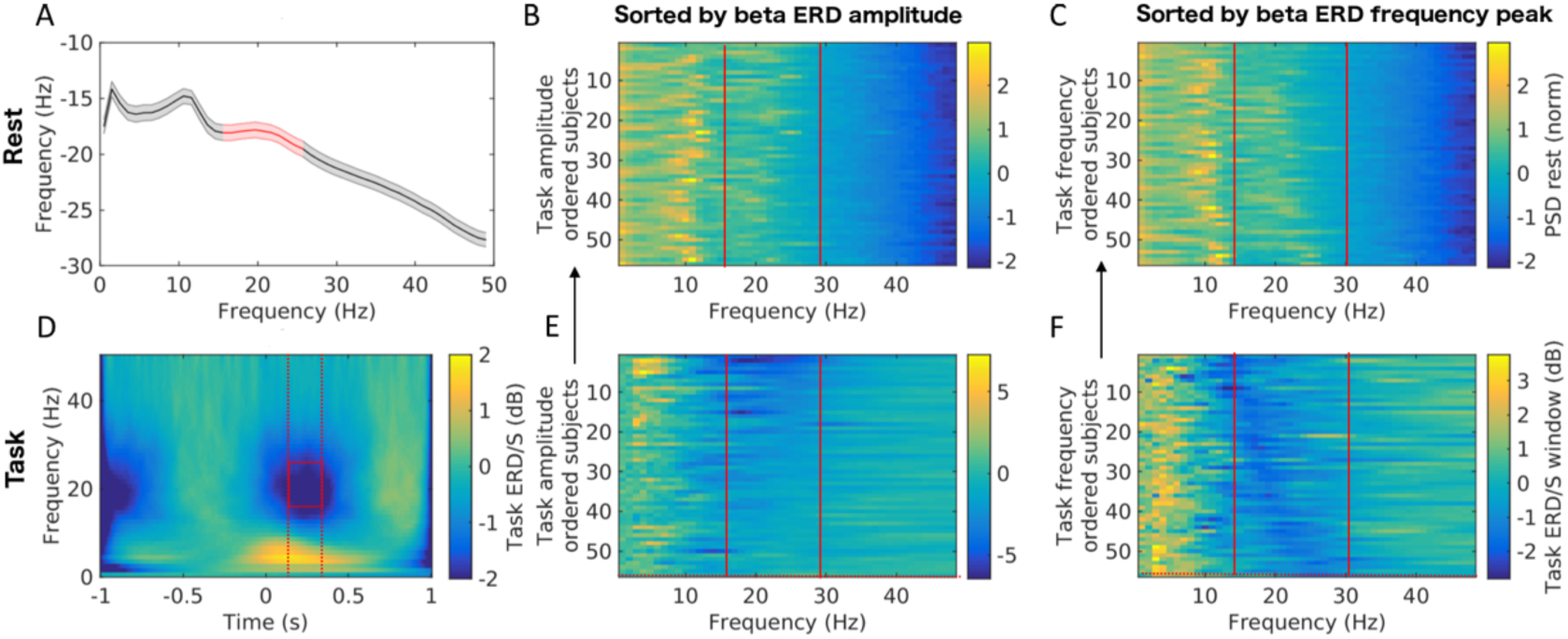
Illustration of the subject variability in task and rest spectral content, and the potential relationships between them. Here, we use the right-hand movement task, locked to EMG onset, as an example. Note that the top row shows rest data, and the bottom row shows task data. **A**. Group-averaged power spectrum of resting data from a parcel in the left motor cortex (data was high-pass filtered above 1Hz). **B, C**. Subject-wise representation of resting state spectra ordered by task features – see below in E and F for type of sorting. **D.** Time-frequency amplitude response in the task condition, showing a beta-band event-related desynchronisation (ERD) (red square). **E.** Task power spectra during the ERD for each subject, ordered by their beta ERD amplitude (averaged over the red time-frequency window in D). Note that the task power spectra in E and F were computed by averaging over the ERD time-period, i.e. within the time-window indicated by the dashed red-lines in D. This ordering index was used in B. **F.** Same power spectra as E, but now with subjects ordered by their beta ERD peak-frequency (found within the time-window indicated by the dashed red-lines in D) - this subject ordering index is used for the resting state data in C.

**Figure 3.**
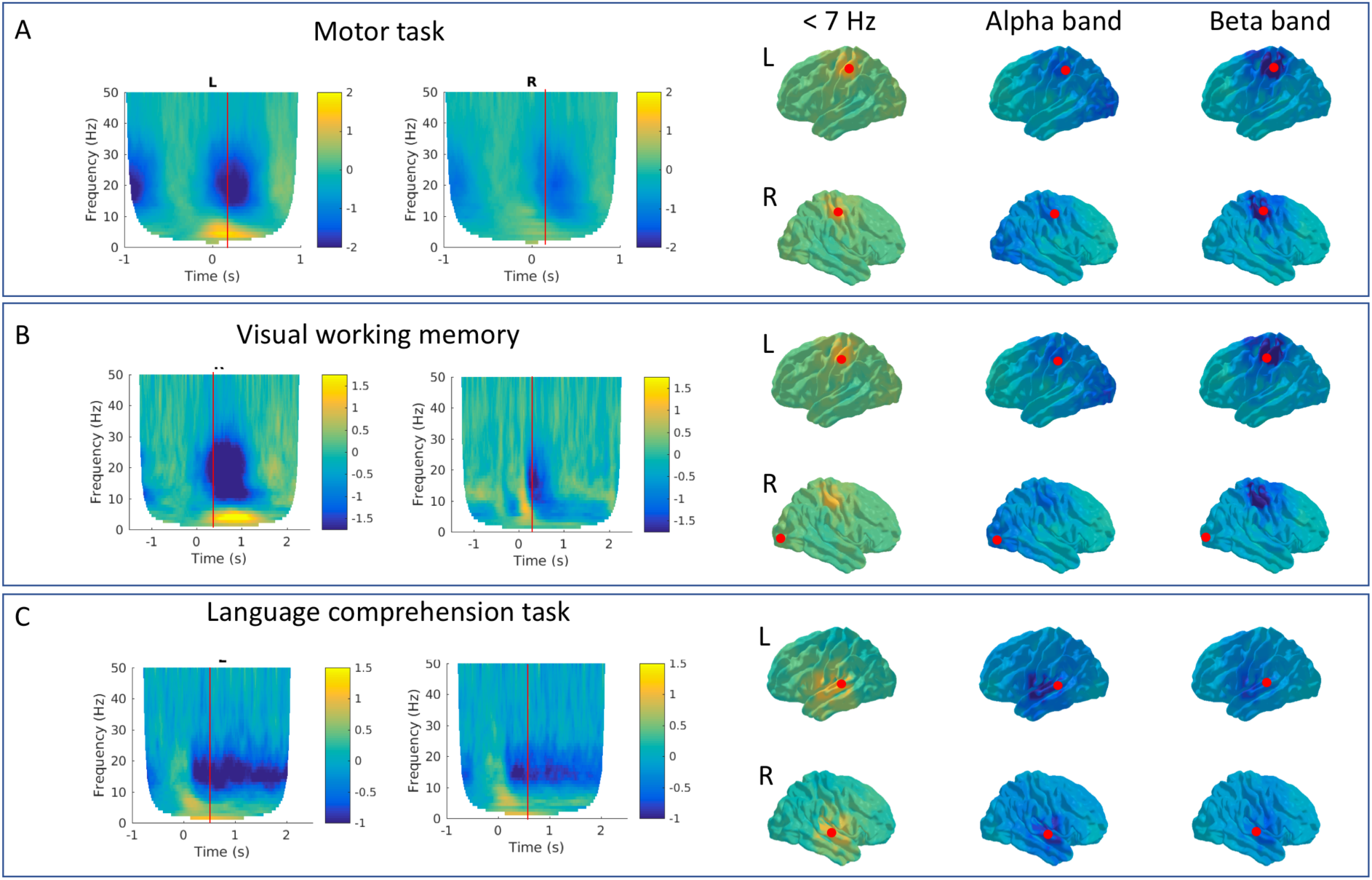
Group-level summary of the different task responses used in this study. **A.** Motor task (a right-hand movement, time-locked to the movement onset). **B.** Visual working memory task (2-back, faces, time-locked to appearance of the non-target items). **C.** Language comprehension task (time-locked to beginning of a sentence). On the left of each panel, is the wavelet-based time-frequency maps locked to task onset, for the parcels indicated by the red dots on the rendered brains on the right of each panel. The red line in the time-frequency plots indicates the time-point show in the rendered brains on the right side, which are shown for three different frequency ranges, corresponding to sub-alpha (<7 Hz, including theta and delta range), alpha, and beta.

**Figure 4.**
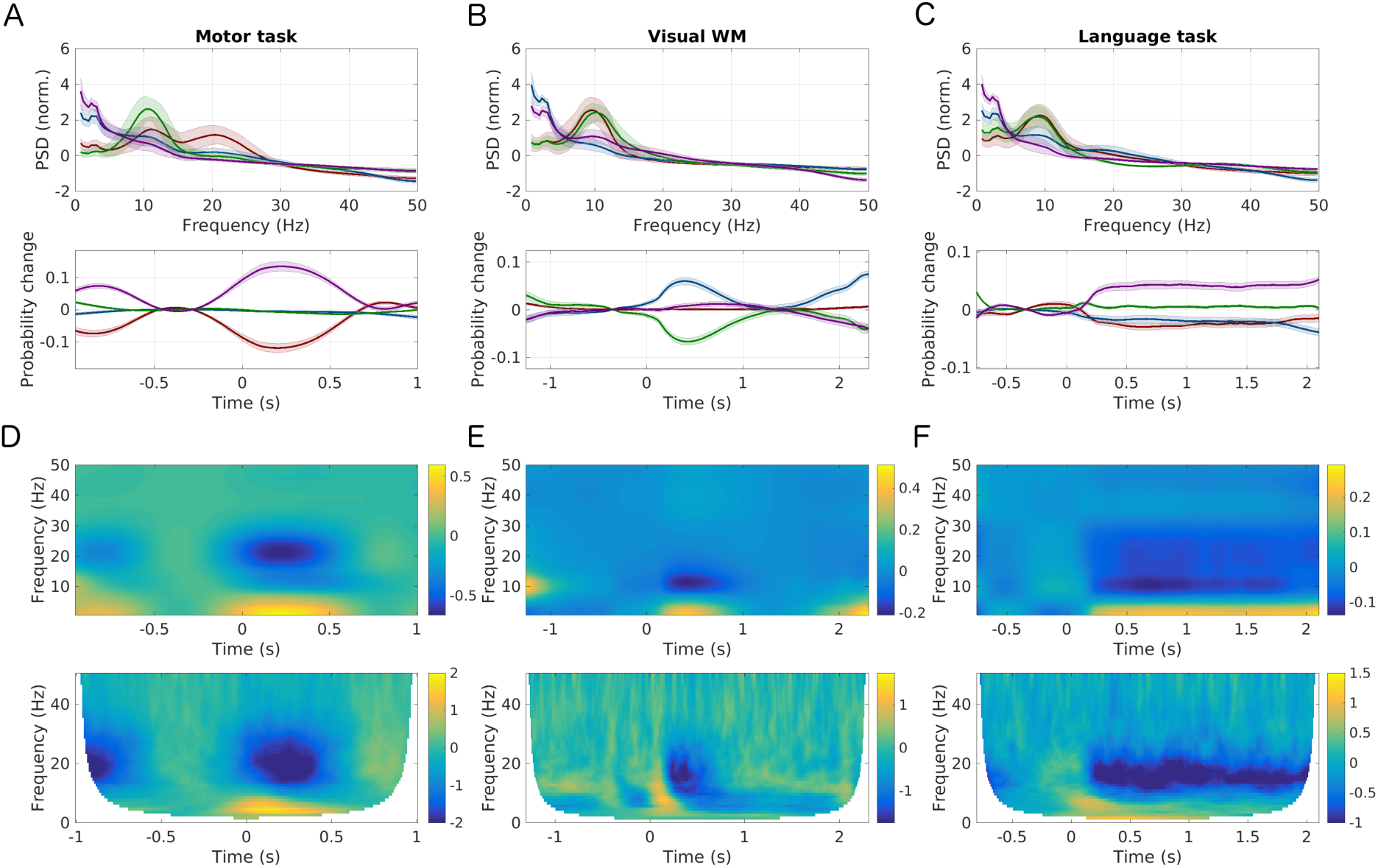
Group result HMM-regularised task responses for all three task groups (motor, WM, language), showing a typical parcel for each condition and its comparison to the conventional wavelet-based time-frequency task maps. **A-C.** HMM state-specific spectral profiles (using state-wise multi-tapering) for the motor task, visual WM task and language task (from different parcels, close to motor cortex, visual cortex and auditory cortex, see inlets). **D-F.** Group average task fractional occupancies (FO), showing the rate of occurrence of each HMM state for the indicated parcel and task. Both the motor task and the visual WM task are reflecting the periodic nature of the experimental design – i.e. preceding or following stimuli can also be seen within the analysed task window (for example, D shows a preceding stimulus, E includes a following one). In F, the language comprehension shows a more sustained response (alpha/beta power decrease, theta-delta increase) corresponding to a less periodic experimental design of that task (one sentences lasts several seconds). Note, that the task fractional occupancies are baseline corrected for visualisation purposes. **G-I.** Group-averaged, HMM-regularized task responses. **K-M.** Group-averaged, conventional wavelet-based responses. Note that the state spectra (i.e. the spectral profiles of the different transient spectral event types) used for generation of the HMM-regularized responses are coming from the resting state data only.

**Figure 5.**
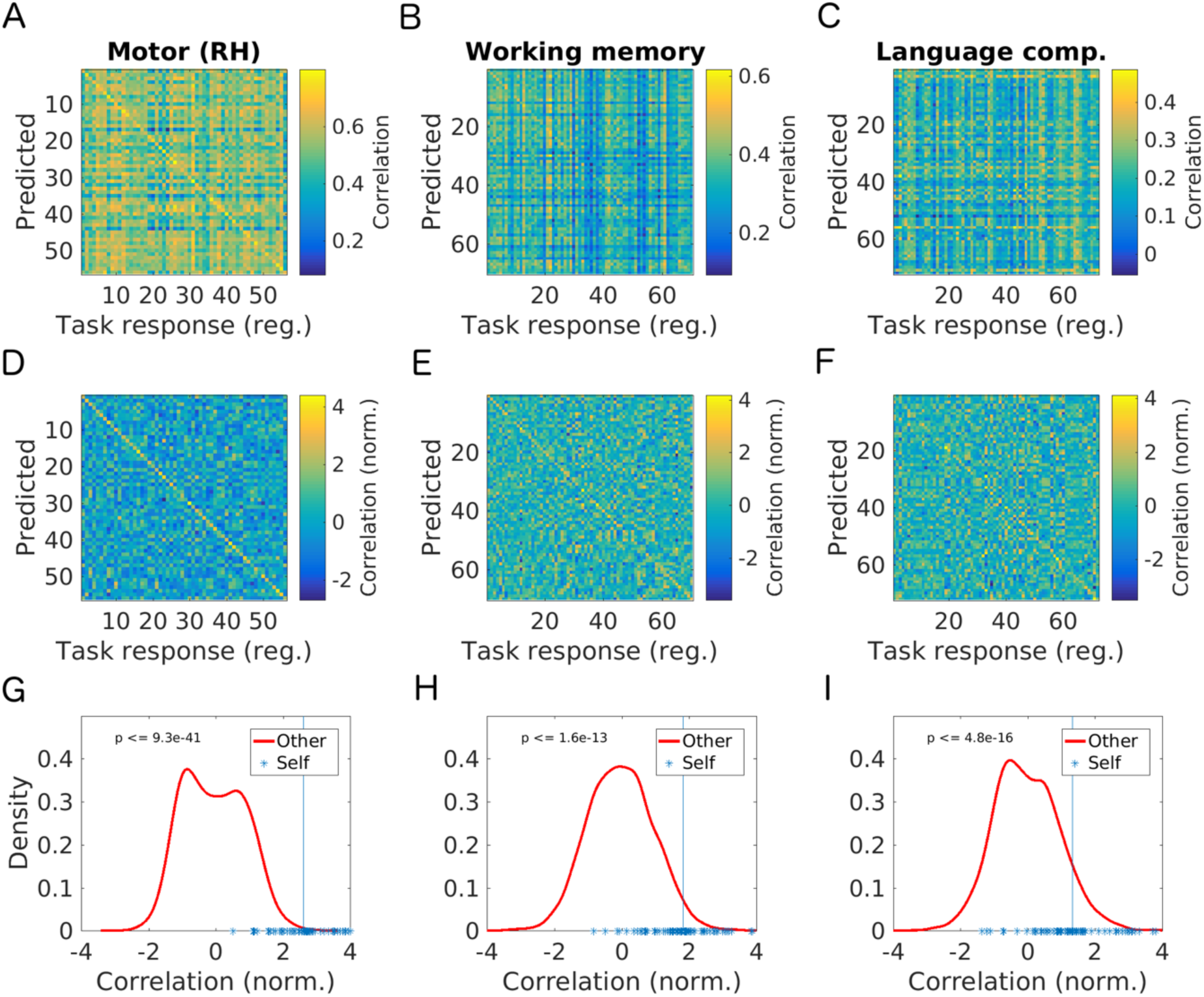
Group level statistics for the prediction results (here one task condition is shown for each group of tasks – motor, working memory and language comprehension - showing the level of correspondence between task responses and predictions. **A-C.** These matrices show the correlation coefficients between actual and predicted task responses of either the same subjects (in the diagonal) or of different subjects (in the off-diagonal). Results are shown for a motor task (right hand movement, A), working memory task (2-back, face stimuli, B) and language comprehension task (sentence understanding, C), respectively. **D-F.** Same task conditions, now with the correlation coefficients normalised over rows and columns (see Methods). **G-I.** Predicted task-responses and actual (HMM-regularized) task responses from the same subjects are more similar to each other (by means of their correlation coefficient, visualised by blue asterisks and the vertical line indicating the mean) than pairs of task response and predictions from different subjects (distribution is visualised by the red lined plot). All task conditions were predicted above chance level.

### 2.6. Influence of genetic factors on prediction of task responses

The aim of the present study is to predict task responses of a subject from their own resting state patterns. Related to this question, we examine whether there is any systematic structure in cross-subject predictions, i.e. when trying to predict one subject’s task response from another subject’s resting state patterns. Specifically, we hypothesize that such a relationship is potentially governed by genetic similarity. The HCP data, specifically the MEG subjects used in this study are selected on grounds of family structure, resulting in roughly a third of monozygotic twins (further denoted MZ), another third of dizygotic twins (DZ) and the remainder being unrelated subjects (UNREL). It has been shown that both for the fMRI and MEG imaging modality, similarity in patterns of functional connectivity in these data sets is related to similarity of genetic structure (Colclough et al., 2017; Vidaurre, Smith, & Woolrich, 2018). We examine whether these different levels of genetic similarity show differences with respect to the capability of our model to predict these subject’s task responses (for example from one twin to another etc.). To do so, after grouping the subjects into groups MZ, DZ and UNREL, for all tasks at hand (n=10) we accumulate the previously computed and normalized correlation coefficients for predicted vs. actual task responses for each pool of subjects (see “Validation of prediction approach” for the normalization approach).

In order to test whether the difference between cross-subject predictability is systematic, we perform permutation tests for the following groups: MZ vs DZ, DZ vs UNREL, with another reference group, SAME, testing for how well we predict from the actual subject (see “Validation of prediction approach”). For each grouping, labels of the two conditions are shuffled 1000 times and we compare the actual group mean difference against the distribution of differences from the permutation distribution.

## 3. Results

### 3.1.1. Data: Subject variability

We start by illustrating the sort of subject variability apparent in task and rest data, and the potential relationships between them. This is shown qualitatively in **Fig. 2,** using the right-hand movement task, locked to the EMG onset. **Fig. 2A** shows the group-averaged resting state spectra; and **Fig. 2D** shows a prominent, typical task induced beta event-related desynchronization (ERD). Next, we looked to see if there were any indications of subject-specific relationships between the spectral properties of the trial-averaged task beta ERD and the spectra in the resting state data.

First, we looked at the **amplitude** of the beta ERD, by plotting in the second column of **Fig. 2** the power spectra in rest and task (calculated during the ERD) over subjects, with the subjects ordered by their task beta ERD amplitude. Second, we looked at the **peak-frequency** of the beta ERD, by plotting in the third column of **Fig. 2** the same power spectra in rest and task over subjects, but now with the subjects ordered by their task beta ERD peak-frequency.

This illustrates two points. First, the subject-specific trial-averaged task and rest spectra show a considerable amount of between-subject variability, in terms of both the amplitude and shape of the spectral profiles. Second, there appears to be a qualitative relationship between the task and rest spectral profiles of individual subjects. Most notably, the task beta ERD **peak-frequency** ordering reveals a similar trend over the subjects between task (**Fig. 2F**) and rest (**Fig. 2C**). For example, subjects that have a high task beta ERD peak-frequency, also tend to have a higher amount of power in high beta than low beta in the rest data. It is these types of relationships that our approach can potentially leverage to allow the prediction of trial-averaged task spectral responses from rest data.

### 3.1.2 Data: Group level task responses

An overview over the typical MEG WL based task responses for the available tasks is given in **Fig. 3**. This shows the group-averaged WL based task responses, i.e. the task-related power changes for each of the three main task conditions – a motor task, visual working memory and a language comprehension task (showing one representative parcel each). The motor task (**Fig. 3A**) shows the typical movement-related alpha and beta ERD in a contralateral motor-cortex associated parcel (approximate parcel location is indicated by red dots on the rendered brains) and a typical motor-evoked response, i.e. a power increase in the lower frequency range (especially in the contralateral motor areas). The group task response for the visual working memory task (**Fig. 3B**) shows both the typical visual alpha ERD following a visual stimulus (which occurs during this task) and the motor preparation component reflected by ERD in the beta band contralateral to the required button press (needed to respond to matches / non-matches). Both the motor component as well as the visual component show power increases in the lower frequency range reflecting the evoked responses usually associated with such a task. The language comprehension task (**Fig. 3C**) shows alpha ERD in a parcel encompassing auditory cortex and higher auditory areas, as well as typical language-related theta power increase. Both are sustained for the duration of the sentence presentation (exceeding beyond the shown time window). These average task responses illustrate that while the tasks share some common spectral features, such as alpha or beta ERD, they vary in their exact spectral profile, temporal dynamics and spatial patterns.

### 3.2. Properties of identified HMM states: spectral features and their modulation in task conditions

We were interested in seeing how well the HMM inferred on resting state data can be used to “predict”, or represent the group-averaged task responses, whether the HMM can capture the task feature on a group level. Note that this *group level prediction* is a pre-requisite to our objective of using the HMM inferred on resting state data to predict *subject-specific* responses. To do this, we compute group-averaged actual task responses, calculated by projecting the HMM state-specific resting power spectra (i.e. the spectral profiles of the transient spectral events represented by each state found in the rest data) onto the task data (see Method section 2.4.4 for computing of the actual task responses). These are then compared with conventional group-averaged WL task responses in **Fig. 4**.

The group-level multi-taper based (MT) spectra for each resting HMM state are shown in **Fig. 4A-C** (for the same parcels chosen as in **Fig. 3**). These are calculated purely from rest data. Then, task HMM state-time courses are calculated by effectively projecting the resting HMM states onto the task data. The resulting group averaged task HMM state occupancies are depicted in 4D-F (these correspond to the rate of occurrence of the different spectral events represented by each HMM state). In all three task groups, these task HMM state-time courses reflect a combination of induced and evoked response dynamics of the task response, showing distinct visible modulations in state probability for each type of dynamics. For example, the purple state in the motor task example (**Fig. 4D**) shows a task-related increase in probability (relative to a pre-movement baseline), with its spectral profile showing a 1/f like behaviour.

By combining the resting HMM state spectral profiles (in **Fig 4A-C**) with the task HMM state spectral profiles (in **Fig 4D-F**) - via means of matrix multiplication – we obtain the actual time-frequency task response for each task, which are shown in **Fig 4G-I**. These can be compared with conventional group-averaged WL task responses shown in **Fig 4K-M**. For example, in the motor task, the actual task response (**Fig 4G**) reproduces low-frequency increases in power, as well as the expected beta ERD known for limb movements. The latter originates from a state with a pronounced beta peak, but which decreases in probability (**Fig 4K**).

Overall, the qualitative correspondence between **Figs. 4G-I** and **4K-M** demonstrates that the spectral properties of the different spectral events (i.e. states) the HMM has identified, from the resting state data, are useful to reproduce qualitative features of the actual task responses of the same group of subjects.

### 3.3. Validation of approach: Group level statistical assessment of single-subject predictions

For the subject-specific predictions we apply the pipeline as outlined in **Fig. 2** (see also Method section 2.4.1: Training and section 2.4.1: Prediction), where we obtain the predicted task responses for all task conditions, parcels and subjects. The predicted task responses are compared to the actual task responses (i.e. the states projected onto the task data, as in **Fig. 4**) by linear correlation analysis. In **Fig. 5**, the validation results for each of the 3 main types of tasks (motor, WM, language) are shown (illustrated for the same tasks as in **Fig 3** and **Fig. 4**). **Fig. 5A** shows the correlation matrix that reflects how strongly the actual task responses correlate with the predictions, either from the same subject (indicated by values in the diagonal) or from other subjects (indicated by values in the off-diagonal part of the matrix). A good prediction should result in the diagonal (prediction of same subjects) prominently standing out. **Fig 5A** shows the ‘raw’ correlation coefficients for this validation step, while **Fig. 5B** shows the same correlation matrix after normalization (of row and columns, respectively). In order to statistically test the difference between same-subject prediction vs random-subject prediction, we tested the null hypothesis that both these groups (same vs other, i.e. in-vs off-diagonal) come from the same distribution (using a two-sample Student’s t-test). This was rejected for all task conditions, as all p-values were less than 1.86 x 10^-5^ (for the math problem solving task). Thus, predictions for all tasks are better when using resting state data (specifically their spectral profiles of the different types of spectral events, or states, as identified by HMM-AR in rest data) from the same subjects as compared to random subjects (defining our ‘chance level’ in the present case).

### 3.4. Single-subject predictions

We next sought to characterise the nature of the between-subject variability that we are predicting. **Supplementary Fig. 2** shows three example subjects, illustrating the between-subject variability that is being predicted in two ways in the right-hand motor task. First, we show the average predicted power in a post-stimulus time-window (190-390 ms) in the beta band. Second, we show the predicted task-related *time-frequency (or spectro-temporal) variability* in a motor parcel contralateral to movement. In summary, this illustrates how the predictions are reflecting both spectro-temporal and spatial aspects of between-subject task variability.

### 3.5 Amplitude and peak-frequency variability drive single subject prediction differentially

After having shown that our approach is capable of predicting task responses both on a group and single-subject level, we wanted to better understand what the model relies on for its predictions, and whether it predicts one particular feature of the task data better than another. In particular, we want to examine whether the model is better suited to predict cross-subject variation in amplitude changes or in other spectral features such as peaks of alpha or beta rhythm ERD.

To this end we first performed principal component analysis (PCA) on a snapshot of the task data. We chose a contra-lateral motor cortical parcel in the right-hand movement task, during a post-movement time window at 190-390ms post-stimulus, using the baseline corrected power over the whole frequency range available (1-50 HZ), averaged over this time window, obtaining one vector of power values per subject. Performing PCA over subjects will reveal the principal components driving subject variability. **Fig. 6** shows the results, focusing on the resulting first (left column) and second principal components (right column) for the task data. The first principal component (PC1) reflects mainly the variation in average amplitude of the beta-band ERD typical for this task, while the second principal component (PC2) mainly reflects subject-specific variations in the peak frequency of this response.

**Figure 6.**
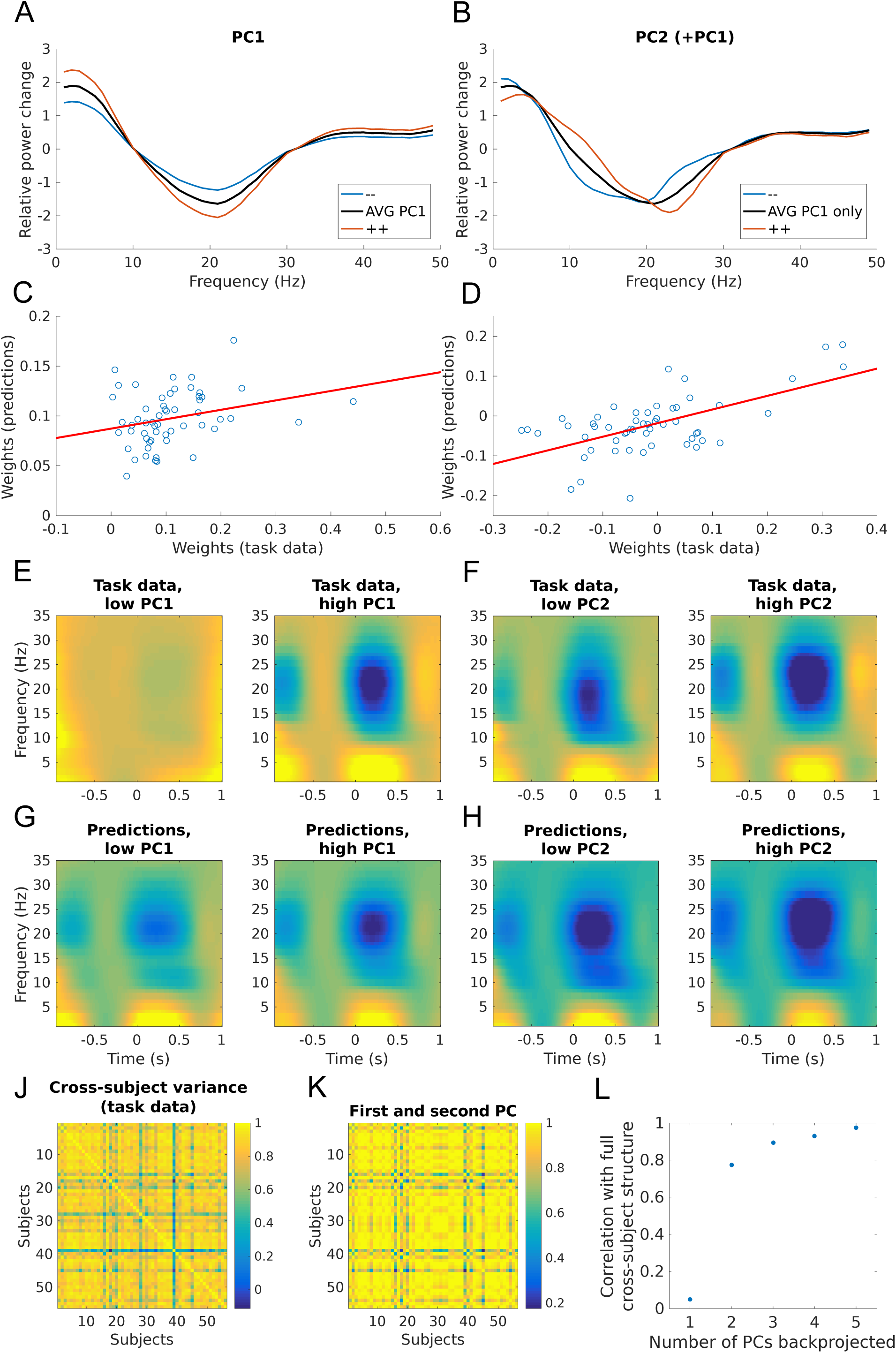
Subject-variability of responses in motor task condition, showing that peak frequency in the beta band is the strongest component of predicted variability. **A,B.** Performing Principal Component Analysis (PCA), on a time-frequency snapshot (over subjects) in a defined post-movement time-window in a motor task on a parcel in contralateral motor cortex A. PCA yields a first principal component (PC) that mainly models the subject-specific mean of the post-stimulus induced (beta-band) power change. The curves labeled with -- and ++ designate lower or higher weightings of PC1 to illustrate its effect (in case of PC1 a simple scaling). B. The second PC appears to model a spectral shift of the beta-band response (higher weights correspond to higher beta frequency). Here, -- and ++ indicate lower and higher weighting of PC2 (added on average PC1 to illustrate its effect). **C,D.** Projecting the predictions onto these components as identified in A and B, the resulting weights across subjects correlate with the actual PCA weights from the task data (rho=0.28 and rho=0.62 for the first and second components respectively). The second PC shows the strongest correlation, indicating a better prediction of spectral features than amplitude features of actual task responses. **E,F.** Performing a median split grouping subjects into low and high weights groups (for each component) shows these amplitude and frequency effects. First component modulates beta ERD while the second one shifts the frequency peak of beta ERD. **G,H.** The same effect is still visible when looking at the projection of the task-PCs onto the predictions. **J,K,L.** We examined the extent to which the cross-subject correlation structure (full in J) is explained by the PCs obtained. **J.** Full cross-subject correlation structure. **K.** By using PC1 and PC2 only (corresponding to beta-amplitude and beta-frequency variations across subjects), back projection into data space results in cross-subject correlations similar to the full data. **L.** Adding PC1 and PC2 for back-projection (n=2) results in correlation of 0.77 with full cross-subject correlation structure. Adding more PCs increases similarity to full structure even more, but not drastically (NB: using the first PC only (n=1) represents a special case, i.e. only modelling the subject-specific mean of beta ERD. Since this does not really yield any relevant cross-subject structure in a correlation analysis, resulting correlation with the full cross-subject correlation matrix is near-zero.

After having run the PCA on the task data, we projected these task-PCs onto the predictions to obtain the weights for this fit (shown in the second column of **Fig 6**). Interestingly, our approach results in PC2 being better preserved (or estimated) in the predictions than the weights from PC1, as is apparent by a higher correlation of the PC2 of the task response with its projection onto the predictions (correlation coefficient r = 0.62 for PC2 compared to r = 0.28 for PC1). Next, the extent to which PC1 and PC2 encodes amplitude or peak variation is demonstrated by sorting the (subject-specific) task responses and the predicted equivalent response according to the weights of PC1 and PC2 (Fig 6C). Overall, **Fig. 6** shows that the model is more capable of reflecting spectral features such as peak variation (e.g. the peak frequency of the beta ERD, as reflected in PC2) rather than amplitude changes (e.g. amount of beta ERD, as reflected in PC1).

In a next analysis, we want to find out how important these two main modes of variation –i.e. amplitude and spectral peak variations - are for explaining cross-subject variability. **Fig. 6J-L** demonstrates that using only these first two principal components, cross-subject variability similar to the ‘full’ task data can be well-represented. This similarity is indicated by a relatively high correspondence of the two matrices that represent the cross-subject variability for the given task and their resulting correlation coefficient of r=0.77. While the correspondence to the full data is even higher for a larger number of retained principal components, the improvement in explained variance is more incremental, and the subsequent components (3 and onwards) are less clear to interpret.

### 3.6. Prediction quality across tasks

Another question that is important to address is the variability of prediction performance, i.e. how well we predict subject specific task responses from rest, in the tasks examined and what is the potential source of this variability in prediction performance. While all tasks yield predictions that are significantly better than chance, the quality of the predictions across task conditions varies to some extent. We hypothesized that one potential source underlying the variability in prediction quality might be the (cross-subject) variability of signal-to-noise ratios (SNR) for the different tasks. The result is visualized in **Fig. 7**. SNR of task responses was defined by the subject’s first-level statistics (i.e. t-values indicating the within-subject level of significance of the observed task responses). It can be seen that for each task - both on a group averaged as well as on the individual subject level - stronger and more stable responses (as indicated by t-values, plotted on the x-axis) imply better prediction (normalized correlation coefficients, plotted on the y-axis). In general, the motor task conditions (right / left hand and feet) are predicted best, followed by the working memory conditions, and finally, the language comprehension task conditions (sentence comprehension, math problem solving). Of interest is that a similar analysis where we examined another potential source of prediction performance variability - effect size, i.e. subject-specific average amount of alpha or beta ERD rather than subject-specific SNR (as done above) - yields no such relationship (results not shown).

**Figure 7.**
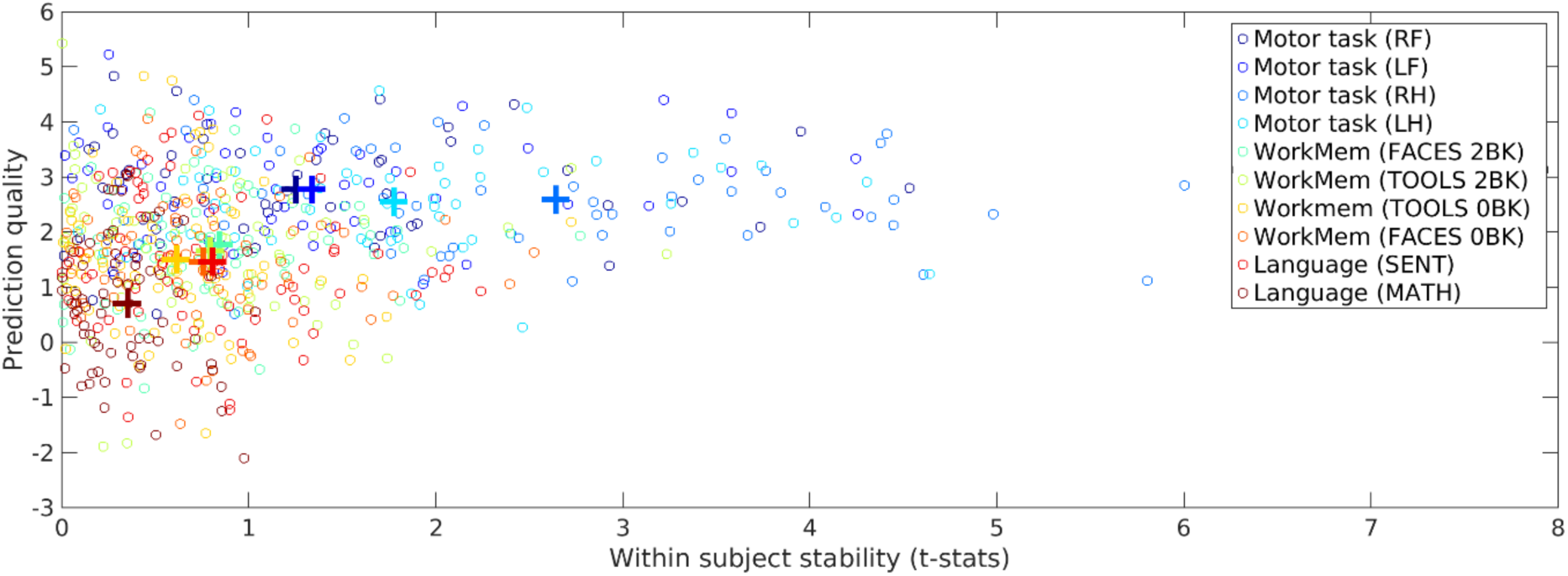
Quality of prediction depends on the subject-specific robustness of actual subject-specific task responses. This scatter-plot visualises the link between first-level significance, or ‘stability’ of task responses (as reflected by the t-values for each subject in a post-stimulus time-window [reflecting the difference from baseline] for a given task condition, plotted on the x axis) and the resulting prediction performance (i.e. the normalised correlation coefficients in the diagonal from Fig. 4, representing the y axis). Average effects across task conditions (as depicted by the coloured crosses) as well as single subjects are shown.

### 3.7. HMM predicted task responses show hereditary structure

We asked how the genetic structure, which is a feature available in the present HCP data set, is related to predictability (**Fig. 8**). We hypothesized that task responses might be better predicted from resting state data of genetically more closely related subjects (e.g. identical or non-identical twins) than task responses predicted from completely unrelated subjects.

**Figure 8.**
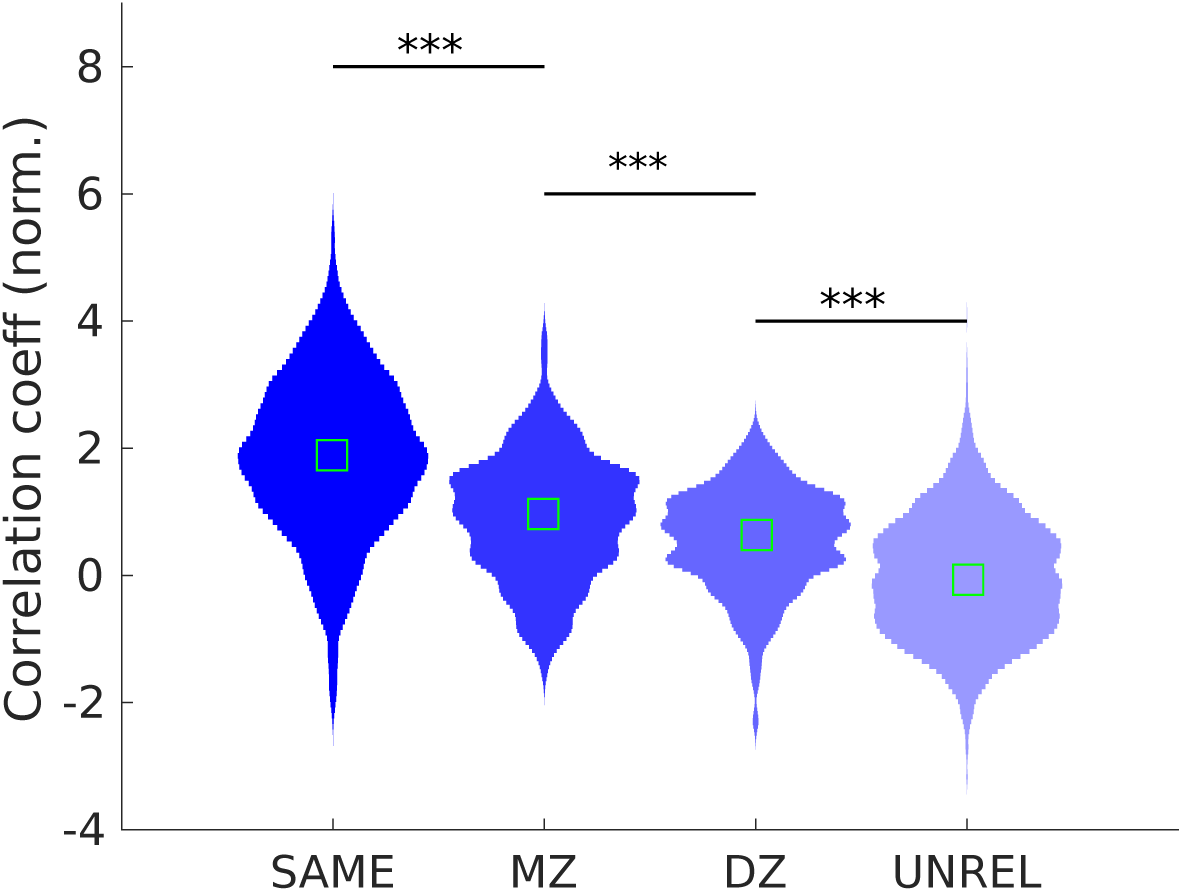
Genetic factors play a role in cross-subject predictions, with subjects being better predicted by their genetically closer counterparts than by unrelated subjects. As expected, when pooling over all same-subjects (‘SAME’), the normalized correlation coefficients are highest (corresponding to the distribution of all pooled diagonal entries from correlation matrices in **Fig. 5**). The second-best prediction across subjects is obtained when predicting task responses from one monozygotic twin’s rest data to its sibling (labeled ‘MZ’, sharing 100% of their genetic information). For dizygotic twins (labeled ‘DZ’, 50% of shared genetic information) predictions are slightly worse (not significant compared to MZ), however they are still significantly better than when predicting task responses of random subjects (‘UNREL’, i.e. unrelated subjects with no shared genetic information). Stars indicate level of significance (***p<0.005, results of permutation testing (with 1000 permutations), following Bonferroni correction for multiple comparisons), light green boxes indicate the median.

Non-parametric analysis of variance revealed that the normalized correlation coefficients (estimating the similarity between prediction and actual task responses, see Methods) pooled into groups; reflecting that same subjects (SAME), identical twins (MZ), non-identical twins (DZ) and the remainder (UNREL) are not originating from the same distribution (as determined by permutation testing). Non-parametric (rank-based) post-hoc testing between the groups showed that all differences in prediction performance between groups were significant (p_SAME_vs_MZ_, p_MZ_vs_UNREL_, p_DZ_vs_UNREL_ all < 0.001, cf. **Fig. 8**). Generally, correlation coefficients are higher (i.e. predictions are better) the more genetically similar subjects are, in terms of the ability for the resting state data from one subject to be able to predict another subject’s task response.

## 4. Discussion

### 4.1. Summary and interpretation of results

We have shown that MEG trial-averaged subject-specific task-activity can be predicted using subject-specific transient spectral events identified in resting-state MEG data using Hidden Markov Modelling. This prediction is made without prior knowledge about the task responses of specific subjects, and is mediated by hereditary factors. This has been demonstrated in a large, freely available data set with a set of diverse experimental conditions ranging from simple hand movements to more cognitively demanding tasks involving working memory, attention and language processing.

The HMM methodology applied here has been previously employed to identify transient events in both rest and task data in an unsupervised manner, and both for electrophysiological as well as hemodynamic data (Baker et al., 2014; Vidaurre et al., 2016; 2018). Here we used a region-by-region HMM-AR to identify transient spectral events defined as having distinct spectral profiles, in order to link rest and task with the following findings.

First, the spectral properties of the different transient spectral events represented by the HMM states extracted from rest were shown to be relevant and effective in describing task dynamics at the group-averaged level. Subsequently, subject-specific spectral profiles of transient spectral events identified by HMM-AR were then used to predict trial-averaged task dynamics on a single-subject level (**Fig. 1**, **Supplementary Fig. 1**). We found that the accuracy of the presented approach typically depends on two factors: The accuracy of the individual spectral profiles of the spectral events as extracted by HMM-AR, and the accuracy of the predicted state dynamics, i.e. the state time-courses or rate of occurrence of the spectral events represented by each state, for the states associated with these spectral profiles. Both these properties critically affect the prediction accuracy, since these two features ultimately generate the predicted (‘HMM-regularised’) time-frequency task responses in our model.

In terms of the general prediction performance, the model tended to predict individual differences in *spectral* features better than differences in the response *amplitude*, as indicated by the results from our principal component analysis of actual task responses and their equivalent counterparts in the predicted responses (**Fig. 6**). In how far this observation is caused by the specific approach chosen (i.e. the HMM-AR), is not completely clear and further work would be needed to elucidate this. In any case, the fact that cross-subject variability of spectral features (especially peak frequency, as encoded in the second PC in **Fig. 6**) is driving a good part of the task prediction implies that these modes might be useful for aligning subjects frequency-wise in the context of task-related analyses, in analogy to the more spatially oriented alignment procedures for fMRI.

With respect to the different task conditions, we saw that task response predictions worked best in task conditions with strong and robust event-related task responses both at the subject and group level (**Fig. 7**). This relationship is likely to come from two different sources. First, a robust task response means that during the training step in our framework, the actual task dynamics, i.e. the state time courses of the hidden states (which serve as a model for the left-out subject) are better estimated when task responses are robust (i.e. less noisy and more genuinely subject-specific); this is true for both subject and group level estimations. Second, a robust response also means that ERD behavior is better estimated in terms of its precise spectral properties (i.e. its peak frequency).

### 4.2. Previous work and related approaches

The results in this work may be relevant for the incipient debate on the interpretation of frequency-specific patterns of neural activity. The success of using the HMM to identify transient spectral events in order to link rest and task speaks in favour for the existence of transient bursts, or a hybrid combination of bursts and sustained rhythms rather than for pure rhythmically sustained oscillations (Shin et al., 2017, van Ede et al., 2018); or at least that there is utility of identifying fast (event-like) transient changes in spectral dynamics. However, a complete comparative analysis with non-bursting representations of spectral activity is required to better support this idea.

In a broader sense, the present study is part of a body of work that tries to link rest to task features, or, more generally, structure to function. Previous approaches have shown links between resting state connectivity patterns and task activations (Biswal, Yetkin, Haughton, & Hyde, 1995; Cole, Ito, Bassett, & Schultz, 2016; Tavor et al., 2016), connectivity and subjects or behavioral measures (Shen et al., 2017; Smith et al., 2015), anatomically related structural features such as grey matter volume linked to behavioral skills such as navigation (Maguire et al., 2000) as well as links between structure and spatio-spectral content (Abeysuriya et al., 2018; Hadida, Sotiropoulos, Abeysuriya, Woolrich, & Jbabdi, 2018). All of these findings point to the functional relevance of inter-subject variance – variance that is necessarily eliminated by conventional approaches such as averaging (Seghier & Price, 2018). This approach builds upon the previous work to show the relevance of this variability in the time-frequency domain by both representing inter-subject variance and showing its link with the resting state.

The evidence that there is a hereditary component (**Fig 10**) - meaning that prediction from resting state data of genetically related subjects yields better task predictions than predicting from unrelated subjects - adds weight to this finding. It supports the idea that these inter-subject differences are not trivial, but biologically meaningful, since related subjects show related MEG patterns. Previous reports already indicated that inter-subject variability, specifically of functional connectivity in resting state (Colclough et al., 2017) as well as spontaneous HMM state (and meta-state) dynamics (Vidaurre et al., 2018) have a strong genetic component, and the finding of genetic influence in the present results is another hint at the relevance of genetic factors for subject variability. Both the spectral profiles and the mapping from resting state to task state dynamics should have some hereditary component to them to yield the present results (being a combination of these two) - however, no systematic comparison has been done to assess their relative contribution more precisely. Regarding the origin of this genetic component – one possibility might be for example cortical folding, which is known to be hereditary to a certain degree – which could affect measurements on the scalp and ultimately source reconstructed rest and task signatures. However, while this might explain spatial variability it is less clear how this would explain spectral variability (such as differences in peak frequency for alpha or beta rhythms, for example).

With respect to predicting task activations from rest, what differentiates our approach from most of the previous approaches (Shen et al., 2017; Tavor et al., 2016) is the challenge of an effectively 3-dimensional task-structure (time-frequency-space), which is more complex than the prediction of static, spatial activation maps (being effectively reducible to 1D representation). Taking this into account, the results are quite encouraging and in terms of statistical robustness comparable to previously reported results for prediction of spatial activation maps in fMRI (Tavor et al., 2016). The data set used in the present study shares a subset of task conditions with the fMRI study (both data sets being part of the Human Connectome Project, (Larson-Prior et al., 2013)) and a subset of subjects. However, apart from the different imaging modality examined – fMRI vs MEG - there is one other noteworthy difference: While our approach seemed to be best at predicting the more ‘simple’ tasks (involving hand or feet movements), and less good at predicting more cognitive tasks (involving subtle changes in relative power), the approach presented in Tavor et al., (2016) showed an opposite effect, performing best in highly cognitive tasks (e.g. language processing). One reason for this might be the sensitivity of our model to spontaneous or induced oscillations such as alpha or beta rhythms prominent in M/EEG resting state – rhythms that are known to be detected (at rest) most clearly in functionally more ‘fundamental’ sensorimotor and posterior visual areas, and detected less in frontal or other more ‘cognitive’ areas higher up in the neural processing hierarchy (Srinivasan et al., 2006).

In a narrower sense, the presented framework, i.e. using HMM-AR and regression-based modeling to predict subject-specific task responses via the identification of transient spectral events, is (in principle) not the only way to predict trial-averaged task responses in M/EEG, or related electrophysiological, data. As outlined in **Fig. 1**, any approach that is capable of decomposing resting state activity into spatio-spectral modes might be similarly used to identify links between rest and task activity. For example, methods like non-negative matrix factorization (Lee, Hashimoto, Wible, & Yoo, 2011), autoregressive modes (Porcaro, Zappasodi, Rossini, & Tecchio, 2009) or other sliding window approaches (O’Neill et al., 2017) are possible options. However, the need for sliding windows - or entirely collapsing spectral features over time - differentiates these from the HMM approach. The unsupervised decomposition of a time-series into consistently reoccurring, transient spectral events with distinct spectral modes - without the need of fixing window length or imposing another temporal structure - is potentially beneficial for the identification of relevant, yet transient and dynamics patterns needed to predict task responses in M/EEG.

Of course, prediction approaches are not at all limited to spatio-spectral features only, but can capitalize on very different properties, for example on static functional connectivity as shown repeatedly for hemodynamic activation maps (Tavor et al., 2016), (Shen et al., 2017)). However, it is not clear whether such an approach would be capable of dealing with the complex spatio-spectral dynamics as it is necessary in the domain of M/EEG time-frequency task responses.

### 4.3. Limitations and challenges

The approach presented here has its limitations. One potential limitation is the assumption of constant spectral features (i.e. stable peak frequency in both rest and task within an HMM state). This assumption may not be always valid. There have been reports of task-induced changes not just of the *amplitude* of ongoing oscillations – which is well established since the discovery of alpha oscillations (Berger, 1929) and later for beta oscillations as well (Pfurtscheller & Lopes da Silva, 1999) – but also changes of the *precise peak frequency* of the induced changes (e.g. shift in the alpha frequency range during increased cognitive demand (Haegens, Cousijn, Wallis, Harrison, & Nobre, 2014). It would be interesting to see whether the incorporation of potential and systematic frequency changes induced by task processing improves prediction quality.

Furthermore, the presented HMM-AR approach used here for prediction of subject-specific task-responses – while operating on whole-brain data via iterating over all available brain parcels – is at its core a mass-univariate approach, since it uses a different HMM-AR on the single time-course of data from each brain region. Thus, by necessity, it ignores any cross-regional interactions. The univariate approach might also be a factor that explains that in our model, sensorimotor task might be predicted better than more cognitive tasks, but this would need further investigation. For future studies, multivariate approaches might be exploited to enable the incorporation of additional features (for example, connectivity measures) with the hope to potentially increase prediction quality. Nonetheless, the mass-univariate approach pursued here has demonstrated success in predicting subject-specific properties in diverse MEG task responses. Utilizing similar HMM methodology to identify transient spectral events in task data, a recent study has shown changes in HMM state dynamics reflecting modulated beta oscillations as a function of motor learning (Zich et al., 2018), adding support to the usefulness and sensitivity of using the HMM with activity from one brain region at a time.

One other important *subject-specific* feature of electrophysiological task responses is largely missing - the latency of evoked and / or induced components - a feature of task responses often used for clinical diagnosis, and a feature reflecting conduction velocity (Halliday, McDonald, & Mushin, 1972), the degree of impairment of fibre tracts, pathways or the impairment of brain areas critically involved in a task-active network. While our model is capable of learning amplitude variations of oscillatory activity to a certain degree, it cannot really predict latency variations in task responses. It is possible for the approach we have developed to be adapted so that *subject-specific* differences in the latency of responses can be taken into account. However, this is not trivial, since the relationship between resting state features and the latency of the task responses is unclear, and thus needs further investigation.

### 4.4. Implications for the functional significance of ongoing activity

One interesting insight with respect to the identified hidden states in our study is that (at least for the given parametrization of the HMM-AR approach chosen here) the approach often appears to assign alpha or beta oscillations to discrete states (**Fig. 4**). Both types of oscillations are known to have functional relevance, modulating task-related activity (Becker, Ritter, & Villringer, 2008; Mazaheri & Jensen, 2008; Nikulin et al., 2007) as well as behaviour (Busch, Dubois, & VanRullen, 2009; Mathewson, Gratton, Fabiani, Beck, & Ro, 2009). This might suggest that these two components do not co-occur, but can be modelled separately - this however is beyond the scope of this study. Interestingly, these oscillatory components are also the aspects of the task response variability that the model predicts best, i.e. in task conditions that have pronounced and stable induced oscillatory changes (specifically often power decreases) in alpha or beta power.

Apart from alpha or beta components, another component that gets well-identified as a discrete state, is a scale-free, broad-band, or 1/f component (see for example the group results for the spectral modes in **Fig. 4**). This type of scale-free activity, reflecting self-similarity or fractality, is thought to have similar functional significance as the more established oscillatory components in M/EEG (He, 2014). In our model, this type of state is relevant and necessary for modelling and predicting the evoked responses in the task data (usually showing a power increase < 7 Hz, as is apparent in **Fig. 3 and Fig. 4**). However, while this 1/f state is the identified HMM-state best suited to modelling these evoked responses, the relationship between this scale-free component in rest and the evoked responses in task is less clear. Here, a higher AR model order might distinguish better between a pure slope and for example, evoked theta to elucidate this question. At the current state, predicting the lower-frequency, evoked components during task processing seems to be less good than modelling and prediction of the oscillatory, induced components. It would have been interesting to establish a closer link between this 1/f type of state in rest and task, however, this is beyond the scope of the present paper.

### 4.5. Outlook & conclusion

Being able to predict time-frequency task responses and their variability in human subjects based on resting state data is highly attractive. Potentially, this might be very useful in a clinical setting. For fMRI, the potential clinical usefulness of task-free neuroimaging has already been demonstrated by predicting the location of language-relevant areas in patients from rest and its feasibility for pre-surgical planning (Parker Jones, Voets, Adcock, Stacey, & Jbabdi, 2017). This suggests that something similar might be achieved with M/EEG in a clinical context. The dominating device in clinical settings is the EEG (and sensor-based analysis), but a similar approach as the one presented here for MEG source data should be feasible as well.

In conclusion, we believe that the framework presented here contributes to the goal of task-free neuroimaging and help to better understand, model and predict inter-subject variability of task responses, identifying the hidden states that serve as the building blocks of this variability.

## Acknowledgments

The Wellcome Centre for Integrative Neuroimaging is supported by core funding from the Wellcome Trust (203139/Z/16/Z). MWW’s research is supported by the NIHR Oxford Health Biomedical Research Centre, by the Wellcome Trust (106183/Z/14/Z), and the MRC UK MEG Partnership Grant (MR/K005464/1). Data were provided [in part] by the Human Connectome Project, WU-Minn Consortium (Principal Investigators: David Van Essen and Kamil Ugurbil; 1U54MH091657) funded by the 16 NIH Institutes and Centers that support the NIH Blueprint for Neuroscience Research; and by the McDonnell Center for Systems Neuroscience at Washington University. We would also like to thank Dr. Alexis Hervais-Adelman for helpful comments regarding the manuscript.

## Supplementary Figures

**Supplementary Figure 1.**
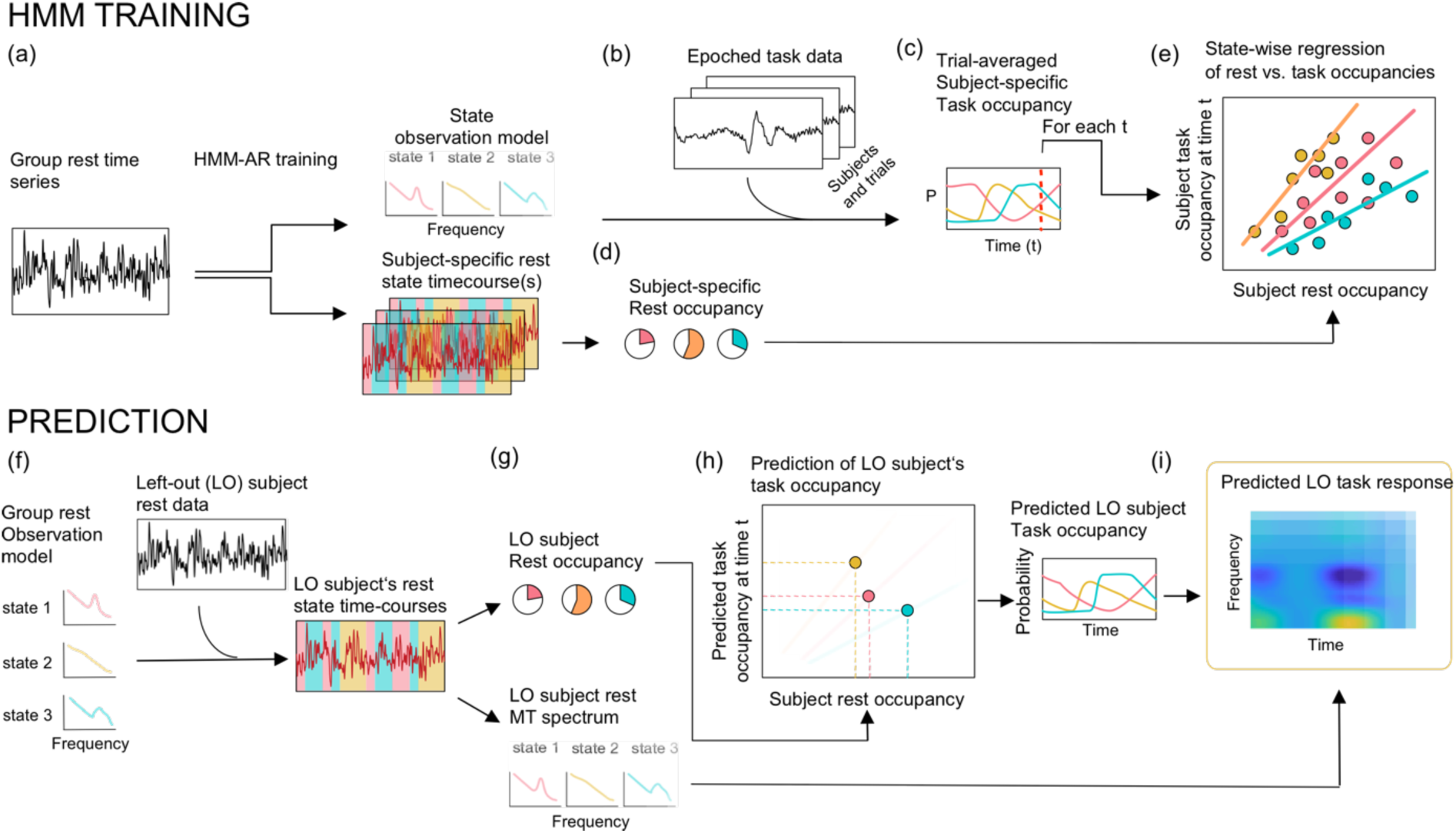
The detailed methodological approach shown for one task condition, and which is carried out separately on all parcels (for ease of illustration, we show only 3 states). **(a-e) Training stage**. (**a**) The group resting state data (one parcel, all subjects concatenated) is fed into an HMM-AR, resulting in the time-courses for each hidden state and subject, and the group-averaged state spectra (i.e. the spectral profiles of the different transient spectral event types) at rest. A hidden state as identified by HMM-AR corresponds to the general concept of a mode as illustrated in Fig. 1. Here, it specifically reflects transient spectral events at rest. (**b**) The HMM-AR is fit to the epoched task data for all subjects and trials, but with the observation models held fixed to the group-averaged state spectra - coming from the transient spectral events at rest - from panel (a). (**c**) The resulting state time-courses are trial-averaged to give the subject-specific task-locked dynamics (or task occupancy) for each state. (**d**) We then do a linear regression across to learn the relationship between the subject-specific rest occupancies (see panel (d)) and task occupancies from (c) at each time-point within a trial. (**f-i) Prediction** of the left-out (LO) subject’s task response. (**f**) The HMM-AR is fit to the LO subject’s rest data, but with the observation model held fixed to the group-averaged state spectra at rest, from panel (a), resulting in the LO subject’s state time-courses at rest. **(g)** The state time courses are used to obtain LO subject state occupancies and its corresponding state spectra that we derived from the transient spectral events at rest. (**h**) The LO subject’s state occupancies in task are predicted by mapping the LO subject’s state occupancies at rest through the linear relationship trained at each trial time-point, from panel (e). (**i**) Finally, the predicted LO subject’s state occupancies in task and the LO subject’s state spectra at rest are combined via matrix multiplication (see Methods) to create a LO subject’s prediction of the time-frequency task response.

**Supplementary Figure 2.**
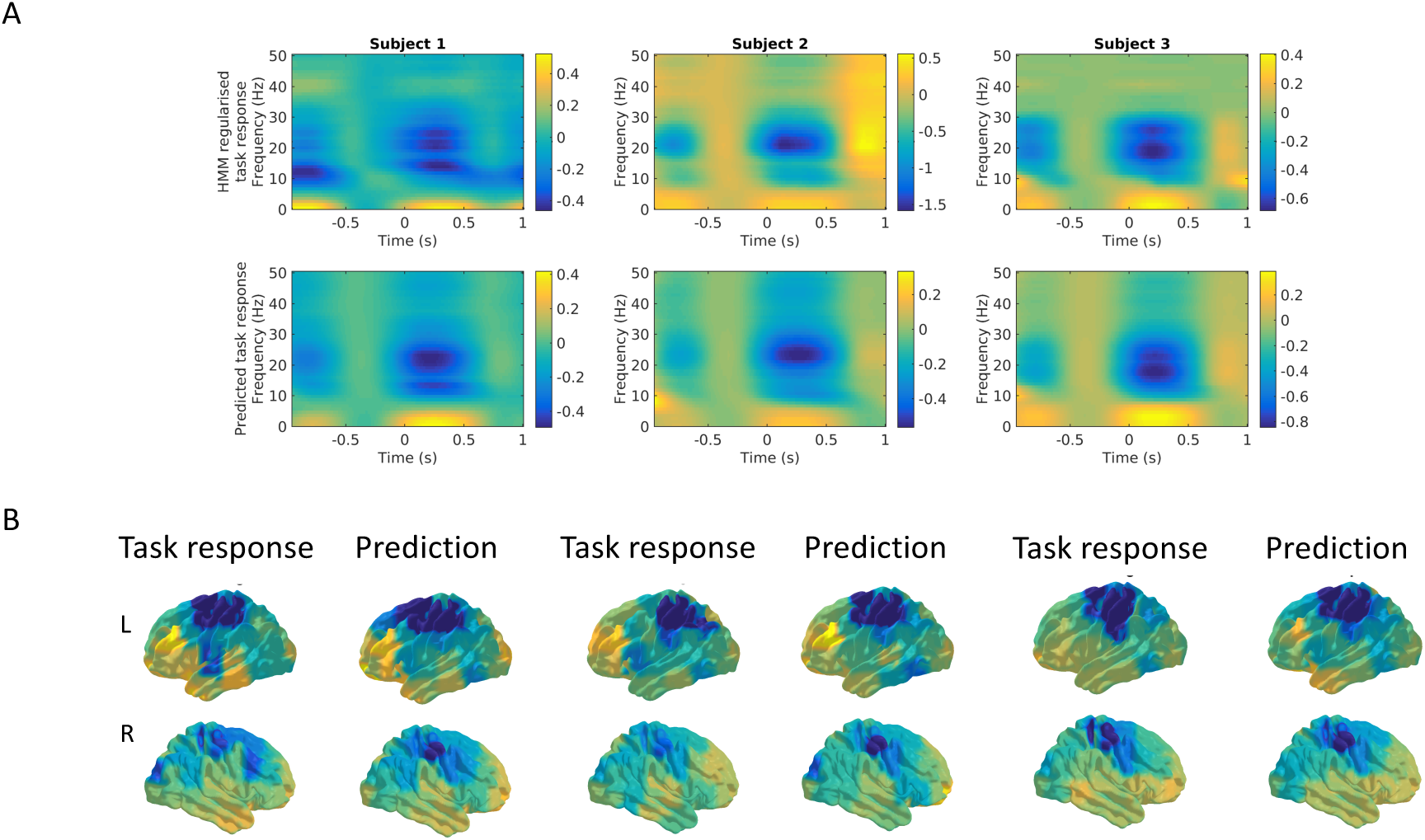
Examples of the between-subject variability being predicted in single-subject task responses, shown for three subjects during the motor task (right-hand movement). **A.** HMM regularized task responses [top] and predicted task time-frequency responses [bottom] (in a motor parcel contra-lateral to the moved hand). **B.** Whole-brain renderings of the induced beta-band response in the actual task data [top] and the predictions [bottom] generated by our approach.

